# Prolactin mediates behavioural rejection responses to avian brood parasitism

**DOI:** 10.1101/2020.10.28.358994

**Authors:** Francisco Ruiz-Raya, Juan Diego Ibáñez-Álamo, Charline Parenteau, Olivier Chastel, Manuel Soler

## Abstract

Even though adaptations resulting from co-evolutionary interactions between avian brood parasites and their hosts have been well studied, the hormonal mechanisms underlying behavioural host defences remain largely unexplored. Prolactin, the main hormone mediating avian parental behaviour, has been hypothesized to play a key role in the orchestration of host responses to brood parasitic eggs. Based on the positive association between plasma prolactin and parental attachment to eggs, decreasing levels of this hormone are expected to facilitate egg-rejection decisions in parasitized clutches. We tested this prediction by implanting Eurasian blackbirds (*Turdus merula*) females with an inhibitor of prolactin secretion, bromocriptine mesylate, to experimentally low their prolactin levels. We found that bromocriptine-treated females rejected mimetic model eggs at higher rates than placebo-treated individuals. To our knowledge, this is the first experimental evidence that host responses to brood parasitism are mediated by the primary endocrine pathway that orchestrates the expression of avian parental care.

## Introduction

Obligate avian brood parasites lay their eggs in the nest of host species, thus exploiting the parental care that other birds provide to their offspring (Roldán and Soler, 2011). This reproductive strategy imposes severe fitness costs on hosts since the newly-hatched parasitic nestling often kills or outcompetes all host’s offspring (Feeney et al., 2014; Soler, 2014). In turn, hosts have evolved a wide range of behavioural, morphological and life-story adaptations to counteract brood parasitism at all stages of the breeding cycle (Feeney et al., 2014; Soler, 2014; Soler, 2017). Surprisingly, while behavioural, cognitive and conditional aspects of host resistance are well known (Soler, 2017), we still have a very limited understanding of the physiological mechanisms underlying anti-parasitic host responses to avian brood parasitism.

Among anti-parasitic behaviours, the recognition and subsequent rejection of parasitic eggs is the most widespread defence used by hosts to counteract brood parasitism (Davies and Brooke, 1989; Feeney et al., 2014; Soler, 2014). At a mechanistic level, egg rejection involves the selective disruption of typical parental behaviours to actively respond against a specific stimulus from the own clutch (i.e. parasitic eggs), so it has been hypothesized that host responses to parasitic eggs might be mediated by hormonal pathways involved in the regulation of parental decisions (Abolins-Abols and Hauber, 2018; Ruiz-Raya and Soler, 2020). Recent evidence suggests that physiological and behavioural host responses to foreign eggs may be mediated by the neuroendocrine pathways underlying stress physiology (i.e. glucocorticoids (Abolins-Abols and Hauber, 2020; Ruiz-Raya et al., 2018)); however, whether rejection decisions are regulated by the endocrine pathways mediating the expression of avian parental behaviours remains unexplored.

Prolactin, classically known as the “parental hormone” (Buntin, 1996; Sockman et al., 2006), is the primary candidate for orchestrating anti-parasitic defences associated with parental decisions (Abolins-Abols and Hauber, 2018; Ruiz-Raya et al., 2018). This peptide hormone mediates the transition from sexual to parental activity in birds (Buntin, 1996; Smiley, 2019; Sockman et al., 2006), playing a key role in the regulation of major aspects of avian parental care: prolactin levels are positively related with the initiation and maintenance of associative parental behaviours such as incubation or post-hatching parental care (Angelier and Chastel, 2009; Angelier et al., 2016; Smiley, 2019). Importantly, the presence of foreign eggs has been found to lead to changes in the prolactin response to stressors (Ruiz-Raya et al., 2018), which suggests that this hormone might play a crucial role in mediating host responses to parasitic eggs (Abolins-Abols and Hauber, 2018; Ruiz-Raya and Soler, 2020). Based on the positive association between circulating prolactin and the expression of avian parental behaviours (Angelier et al., 2016; Smiley, 2019), it may be hypothesized that a reduction in prolactin levels would affect rejection decisions. Specifically, decreasing prolactin levels would be expected to result in more restrictive egg acceptance thresholds and facilitate rejection decisions (Abolins-Abols and Hauber, 2018). Here, we tested this hypothesis for the first time by implanting host females with bromocriptine-mesylate pellets, a dopamine receptor agonist, to experimentally decrease their prolactin levels. Afterwards, we assessed their response to experimental parasitism compared to placebo-implanted individuals in order to determine whether changes in prolactin profiles affect hosts’ rejection decisions.

## Material and methods

### Study site and species

This study was conducted in a Eurasian blackbird (*Turdus merula*) population located in the Valley of Lecrín, Southern Spain, from late March to June 2018. The Eurasian blackbird (hereafter blackbird) is a potential common cuckoo (*Cuculus canorus*) host frequently used in egg-rejection studies (e.g. Grim et al., 2011; Roncalli et al., 2019; Ruiz-Raya et al., 2015; Samas et al., 2011; Soler et al., 2015; Soler et al., 2017). Blackbirds typically remove foreign eggs by grasping them with their bill (Soler et al., 2015), showing high egg-ejection abilities (see references above).

### Hormone manipulation

Females (N = 36) were captured just after clutch completion (mean ± se = 2.3 ± 0.19 days) by using a mist net placed near the nest (6:00 - 8:00 am). Blood samples were collected from the brachial vein with a 25-gauge needle and 80-μl heparinized microhematocrit tubes within 3 minutes after capture. Through the breeding season, we sequentially assigned females to one of two experimental groups. Half of the females (n = 18) were implanted with bromocriptine mesylate (BRC) time-release pellets (according to the characteristics of our study species: C-231, 0.5 mg, 10-days release, 3.1 mm diameter; Innovative Research of America), a dopamine receptor agonist which has been widely used to lower circulating prolactin in avian species (e.g. Angelier et al., 2006; Cottin et al., 2014; Smiley and Adkins-Regan, 2018; Thierry et al., 2013). Another half of the individuals (n = 18) were assigned to the control group and therefore implanted with placebo pellets (C-111, 0.5 mg, 10-day release, 3.1 mm diameter; same company). Time-release dosage pellets have been frequently used to modify hormone levels in ecophysiological studies, showing no negative long-term effect for birds’ welfare (e.g. Cottin et al., 2014; Müller et al., 2009; Thierry et al., 2013). Even so, in order to discard the existence of unintended side effects of our BRC treatment on avian feeding behaviour (Buntin, 1989), females were weighted to the nearest 1 g before and after the hormonal treatment to assess potential changes in their body mass (Smiley and Adkins-Regan, 2018). Additionally, although nest desertion is not an egg-rejection mechanism in blackbirds (Soler et al., 2015), we evaluated whether our BRC manipulation affected the probability of nest abandonment.

BRC and placebo-implanted individuals underwent identical procedures. Pellets were implanted subcutaneously in the female’s back through a small incision (5 mm) that was subsequently sutured with surgical tissue adhesive (Vetbond, 3M). In our study population, there was no seasonal variation in baseline prolactin in both experimental groups (both cases *p* > 0.57; Fig. S1; see statistical methods). Individuals were marked with a combination of coloured rings to verify their identity. All birds were released 10 min after implantation. All females returned to their nest to resume incubation within the next two hours. To test the effectiveness of the 10-day release BRC pellets, females (N = 9) were recaptured six days after implantation (i.e. at the end of the experimental parasitism trials; see below) and a second blood sample was collected from the opposite wing. Similar sample sizes have been proved valid to verify hormone implants/injection effectiveness in previous studies (Abolins-Abols and Hauber, 2020; Goutte et al., 2010; Ouyang et al., 2013), so we minimized the number of re-sampled females to reduce the stress linked to recapture and handling for breeding pairs. The visual examination of recaptured individuals revealed that incisions were successfully sutured in all cases. Blood samples were kept cold for up to 6 hours until centrifugation. Afterwards, plasma was extracted and stored at −20 °C until the hormonal assay. Prolactin plasma concentration was determined by a heterologous radioimmunoassay at the Centre d’Etudes Biologiques de Chizé. The assay was conducted using chicken prolactin and antibody against chicken prolactin (supplied by Dr. A. F. Parlow, Harbor-UCLA Medical Center, Torrance, CA, USA). All samples were run in one assay, in duplicate, with 25 μl of plasma for each duplicate and a limit of detection of 0.43 ng/ml. The intra-assay coefficient of variation was 10.3%.

### Brood parasitism experiments

Nests were parasitized with mimetic eggs 24 hours after pellet implantation. As model eggs, we used natural blackbird eggs collected from deserted clutches, which were painted mimetic to simulate interspecific parasitism (Fig. 2b; see (Soler et al., 2015) for additional details). This type of mimetic model eggs elicits intermediate ejection rates in blackbirds (26.8 - 50% (Roncalli et al., 2019; Soler et al., 2015)), thus allowing to detect bi-directional changes in egg-rejection rates in response to the experimental manipulation. All nests were checked every 24 h to determine egg ejection. Recent research shows that nest desertion is not an egg-rejection mechanism in medium-sized hosts such as blackbirds (Soler et al. 2015), so nest abandonment was not considered a genuine response to experimental brood parasitism. Experimental eggs were consider as accepted if they remained in active nests for 5 days (Roncalli et al., 2019; Ruiz-Raya et al., 2015; Ruiz-Raya et al., 2016; Soler et al., 2017).

### Statistical analysis

Between-groups differences in prolactin levels prior to our hormonal manipulation were assessed through a linear model including treatment, clutch size, and their interaction as predictors. We used linear regression to explore seasonal variations in baseline prolactin levels before implantation. Linear mixed-effect models were fitted by using *nlme* (R package v.3.1-117 (Pinheiro et al., 2014)) to assess the effect of our manipulation on circulating prolactin and body mass. As fixed factors, our mixed models included: implant treatment (BRC or placebo), sample day, and their interaction, while female ID was included as random term. A generalized linear model with binomial error was performed to assess the effect of our experimental treatment on egg ejection. As predictors, our binomial regression model included: implant treatment, clutch size (3-4 eggs), the interaction between these two terms, baseline prolactin levels before implantation and laying date. We assessed whether our BRC treatment affected nest abandonment through a binomial regression model including implant treatment and clutch size as predictors. Stepwise procedures were avoided to minimize the chances for type I errors, and full models were used for inference (Whittingham et al., 2006). Assumptions for normality of residuals and homogeneity of variances were verified by the visual inspection of the residual graphs. All analysis and graphs were performed using R version 3.6.1.

## Results and Discussion

Pre-treatment prolactin levels did not differ between BRC and placebo-implanted females (estimate ± se = 3.459 ± 6.493; t = 0.533; *p* = 0.60; N = 36; Fig. 1a); however, both groups differentially varied their prolactin levels in response to our BRC treatment (implant treatment x sample day: F_1,7_ = 52.17; *p* < 0.001; Fig. 1b). Specifically, while BRC-treated individuals showed a significant reduction in their prolactin levels after implantation (Tukey’s test: estimate ± se = 35.39 ± 4.02; df = 7; t = 8.81; *p* < 0.001), circulating prolactin did not vary in placebo-implanted females (Tukey’s test: estimate ± se = −8.12 ± 4.49; df = 7; t = −1.81; *p* = 0.34). These results confirm the effectiveness of our hormonal manipulation, and supports previous studies using BCR to experimentally decrease plasma prolactin levels in birds (e.g. Angelier et al., 2006; Cottin et al., 2014; Smiley and Adkins-Regan, 2018; Thierry et al., 2013).

**Figure 1:**
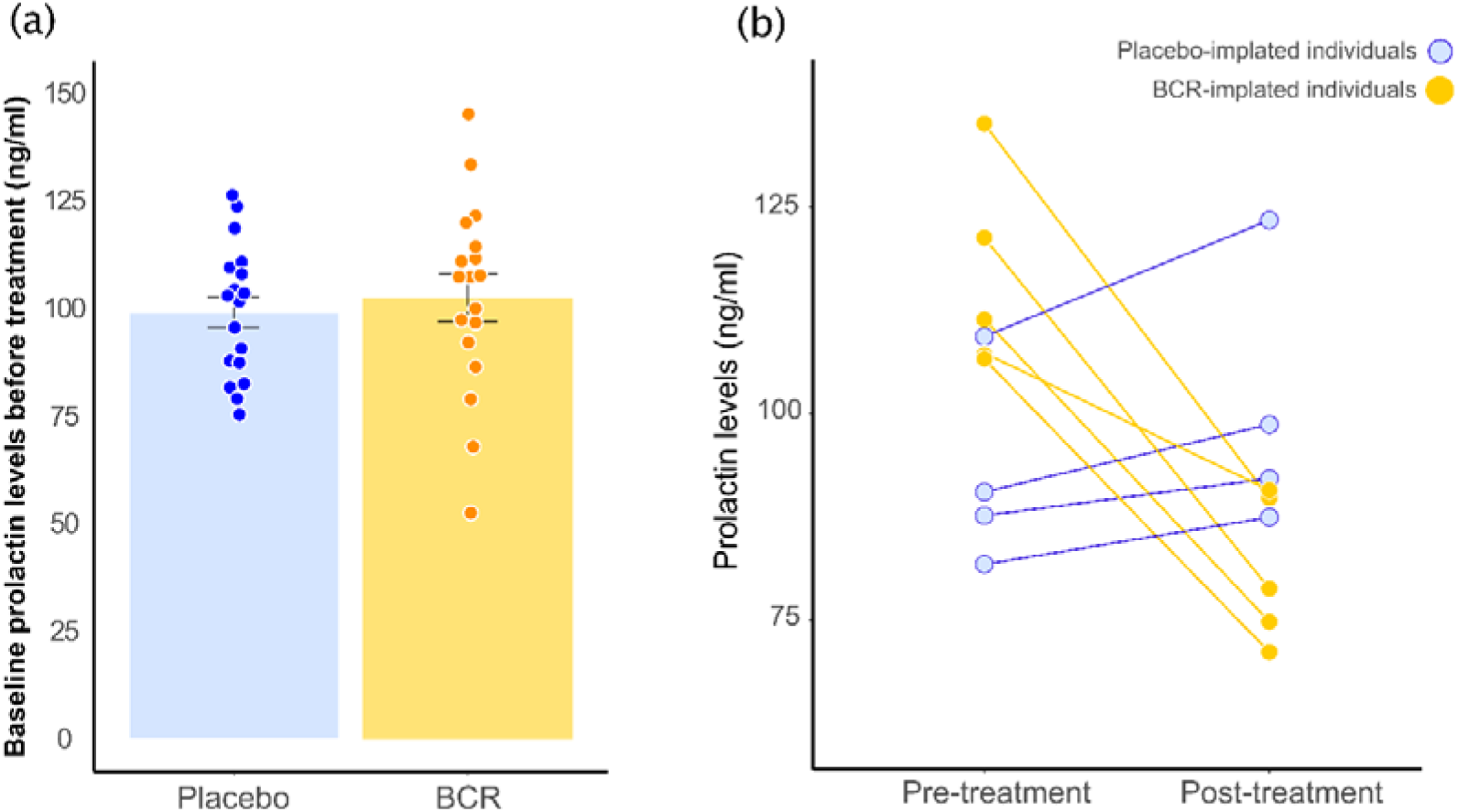
(a) Pre-treatment prolactin levels (i.e. before implantation) showed by placebo (n = 18) and bromocriptine-implanted (BRC; n = 18) females. (b) Effect of the bromocriptine (BRC) 10-days-release pellets on females’ prolactin levels 6-days after implantation: BRC-implanted females showed a significant reduction in circulating prolactin post-treatment compared to placebo-implanted individuals.

Prolactin is the main hormone orchestrating the expression and maintenance of avian parental behaviours during incubation (Angelier and Chastel, 2009; Smiley, 2019), so egg-rejection decisions might be affected by circulating levels of this hormone (Abolins-Abols and Hauber, 2018; Ruiz-Raya and Soler, 2020). As expected, the experimental decrease in circulating prolactin carried out in our study significantly affected the host response to experimental brood parasitism: BRC-implanted females ejected mimetic model eggs at significantly higher rates than placebo-implanted individuals (implant treatment: χ^2^ = 5.59; df = 1; *p* = 0.018; Fig. 2a), independently of clutch size (implant treatment x clutch size: χ^2^ = 0.24; df = 1; *p* = 0.62). Laying date and baseline prolactin levels did not affect the egg-ejection behaviour of females (both cases *p* > 0.76). These results show that the experimental reduction of plasma prolactin increases the probabilities of egg rejection in blackbirds, while placebo-treated individuals ejected mimetic model eggs at similar rates to those found in previous studies in this species where no hormonal manipulation was performed (Soler et al., 2015). To the extent of our knowledge, this is the first experimental demonstration that host responses to brood parasitism may be mediated by the main endocrine pathway involved in the expression of avian parental care.

**Figure 2:**
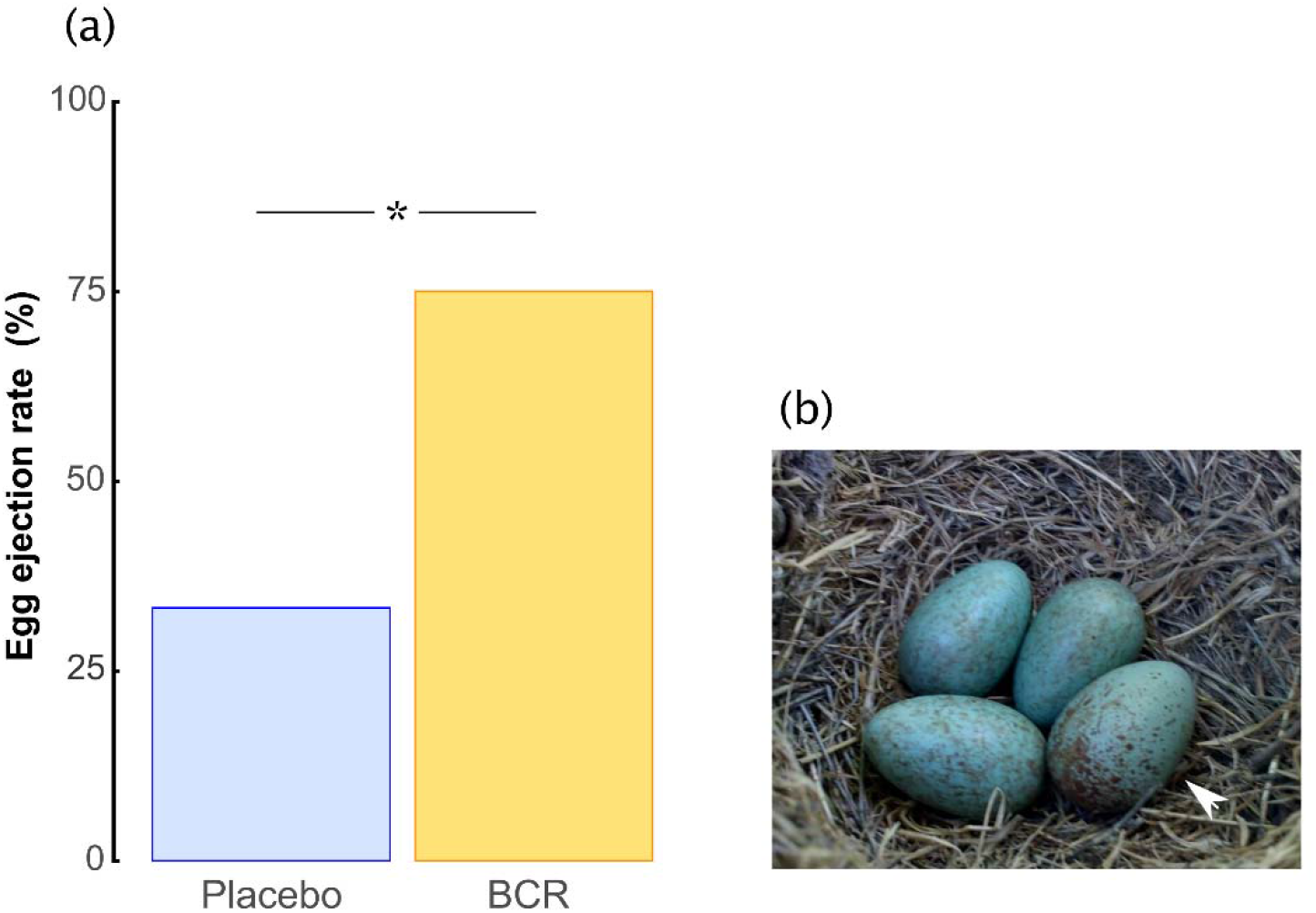
(a) Probability of egg ejection in both bromocriptine (BRC; yellow; N = 16) and placebo-implanted females (Placebo; blue; N = 15). (b) Blackbird clutch experimentally parasitized with a mimetic model egg (white arrow).

A better knowledge of the endocrine bases of host responses to brood parasitism will be crucial to further understand individual variation in the expression and evolution of rejection decisions (Abolins-Abols and Hauber, 2018). Since hosts might mistakenly eject an own egg (recognition costs), or break their clutch during the process (ejection costs) (Marchetti, 1992; Stokke et al., 2002), parasitized individuals are expected to optimize their individual performance by adjusting their acceptance thresholds according to host-parasite egg phenotype dissimilarity, previous experience or parasitism risk (Hauber et al., 2006; Ruiz-Raya and Soler, 2017; Ruiz-Raya and Soler, 2020). While the endocrine mechanisms underlying conditional responses to avian brood parasitism have been little studied, our results indicate that flexible shifts in hosts’ acceptance thresholds may be linked to variations in circulating prolactin.

Interestingly, Abolins-Abols and Hauber (2020) have recently shown that an experimental reduction in plasma corticosterone increases the acceptance of parasitic model eggs in American robins (*Turdus migratorius*), a brown-headed cowbird (*Molothrus ater*) host closely related to Eurasian blackbirds. Taken together, these findings suggest that egg-rejection decisions are under the control of multiple endocrine mechanisms, and draw attention to the need for integrative studies combining prolactin and corticosterone, two hormones that may interact under certain circumstances to maintain parent homeostasis, providing complementary information on parental decisions (Angelier et al., 2016).

Beyond individual differences in baseline prolactin levels, interindividual variation in egg-rejection decisions might be determined by differences in hormonal sensitivity to environmental stimuli. Plasma prolactin has been found to decrease in response to stressors in many bird species (Angelier and Chastel, 2009; Angelier et al., 2016), so the host response to brood parasitism might be to some extend modulated by the strength of the prolactin response to environmental stressors. For example, an increase in the perceived risk of brood parasitism (e.g. witnessing a parasite near the nest) might lead to a drop in host’s prolactin levels, which would explain why the sight of a parasite at the nest facilitates egg rejection by some hosts (Bartol et al., 2002; Moksnes et al., 1993; Moksnes et al., 2000). Under certain circumstances, hosts might be expected to modulate their prolactin response to stressors according to current environmental conditions, which could result in an adaptive prolactin stress response (Angelier and Chastel, 2009). Some hosts rely on the combined use of personal and social information about local parasitism risk to plastically adjust their acceptance thresholds to the current environmental context (Thorogood and Davies, 2016), or they require from two stimuli of brood parasitism (a parasitic egg in the nest and female parasite presence) to reach the threshold for egg rejection (Moksnes et al., 1993). Interestingly, the reduction in plasma prolactin in response to stressors is more pronounced in individuals experimentally parasitized with non-mimetic eggs (Ruiz-Raya et al., 2018), which could suggest that prolactin responsiveness to parasite presence might be higher after perceiving a first cue of brood parasitism (e.g. an odd egg in the nest). While brood parasitism has been shown to impact the endocrine profiles of adult hosts during incubation (Ruiz-Raya et al., 2018), nestling (Antonson et al., 2020) and fledgling stages (Mark and Rubenstein, 2013), the extent to which adult parasite presence affects the hormonal levels of hosts still needs further study. Furthermore, since circulating prolactin often vary according to different life-history contexts (e.g. in response to predation risk or unpredictable environmental conditions (Angelier and Chastel, 2009; Angelier et al., 2016)), our understanding of the physiological mechanisms underlying anti-parasitic defences would benefit from the study of the endocrine response of hosts to their entire environmental context.

In our study, five out of 36 females (13.8%) deserted their nest but, most importantly, this abandonment probability was independent of the hormonal manipulation (χ^2^ = 0.23; df = 1; *p* = 0.63) and clutch size (χ^2^ = 0.07; df = 1; *p* = 0.80). Decreasing prolactin levels is associated with nest abandonment and lower breeding success in some bird species (Angelier et al., 2016), although this link is not clear in other avian species (Angelier et al., 2016; Kosztolányi et al., 2012; Wojczulanis-Jakubas et al., 2013). Our results confirm previous studies showing that nest desertion is not predicted by changes in circulating prolactin in Eurasian blackbirds (Ruiz-Raya et al., 2018), a potential host that does not use nest desertion as an anti-parasitic defence (Soler et al., 2015). These results suggest that nest abandonment is likely under the control of different neuroendocrine pathways, as well as other important aspects of avian breeding biology such as energy storage (Angelier et al., 2016; Kosztolanyi et al., 2012). Future studies should explore whether prolactin changes are linked to nest desertion decisions in small-sized hosts that use nest abandonment as anti-parasitic defence. Our experimental manipulation had no impact on females’ body mass (sample day: F_1,7_ = 1.98; *p* = 0.20) independently on the hormonal treatment (implant treatment x sample day: F_1,7_ = 1.02; *p* = 0.35). These results indicate that our BRC treatment did not effected females’ feeding behaviour (Buntin, 1989), as well as supports previous results showing absence of unintended side effects on feeding behaviour in BRC-treated zebra finches (*Taeniopygia guttata*) (Smiley and Adkins-Regan, 2018).

In conclusion, our findings show that prolactin may play an important role as a mediator of host resistance to avian brood parasitism. While decreasing prolactin levels likely lead to more restrictive acceptance thresholds by facilitating host rejection decisions, the mechanism through which hormonal variations mediate egg-rejection decisions need further exploration. The fact that BRC-treated individuals ejected model eggs at higher rates, as well as the absence of ejection costs, suggests that a reduction in circulating prolactin does not impact the cognitive abilities of hosts. Indeed, changes in the individual cognitive performance are likely mediated by alternative endocrine pathways such as glucocorticoids (Maille and Schradin, 2016). Future studies addressing individual variations in baseline prolactin levels, the strength of the prolactin response to stressors, or differences in target tissue sensitivity to hormones (e.g. prolactin receptor density) would be particularly helpful to further explore the hormonal mechanisms mediating differences in egg-rejection behaviour within and among host populations. Finally, prolactin might also play a decisive role in the regulation of other anti-parasitic host defences linked to parental decisions, such as the rejection of parasitic nestlings (Grim, 2017), so future research on the role of prolactin as mediator of host defences to brood parasitism should be extended to other stages of the breeding cycle.

## Ethics

This study follows all relevant Spanish national (Decreto 105/2011, 19 dee Abril) and regional guidelines. No individual exhibited long-term negative effects as a consequence of our treatment.

## Authors’ contribution

FRR, JDIA and MS designed the study. FRR conducted the fieldwork. CP and OC performed the hormonal assays. FRR analysed the data and wrote the first draft. All authors critically contributed to drafts and gave final approval for publication.

